# Predicting DNA Hybridization Kinetics from Sequence

**DOI:** 10.1101/149427

**Authors:** Jinny X. Zhang, John Z. Fang, Wei Duan, Lucia R. Wu, Angela W. Zhang, Neil Dalchau, Boyan Yordanov, Rasmus Petersen, Andrew Phillips, David Yu Zhang

**Affiliations:** Department of Bioengineering, Rice University, Houston, TX; Systems, Synthetic, and Physical Biology, Rice University, Houston, TX; Microsoft Research, Cambridge, UK

## Abstract

Hybridization is a key molecular process in biology and biotechnology, but to date there is no predictive model for accurately determining hybridization rate constants based on sequence information. To approach this problem systematically, we first performed 210 fluorescence kinetics experiments to observe the hybridization kinetics of 100 different DNA target and probe pairs (subsequences of the *CYCS* and *VEGF* genes) at temperatures ranging from 28 °C to 55 °C. Next, we rationally designed 38 features computable based on sequence, each feature individually correlated with hybridization kinetics. These features are used in our implementation of a weighted neighbor voting (WNV) algorithm, in which the hybridization rate constant of an unknown sequence is predicted based on similarity reactions with known rate constants (a.k.a. labeled instances). Automated feature selection and weighting optimization resulted in a final 6-feature WNV model, which can predict hybridization rate constants of new sequences to within a factor of 2 with ≈74% accuracy and within a factor of 3 with ≈92% accuracy, based on leave-one-out cross-validation. Predictive understanding of hybridization kinetics allows more efficient design of nucleic acid probes, for example in allowing sparse hybrid-capture panels to more quickly and economically enrich desired regions from genomic DNA.

---

Hybridization of complementary DNA and RNA sequences is a fundamental molecular mechanism that underlies both biological processes [1–3] and nucleic acid analytic biotechnologies [4–7]. The thermodynamics of hybridization have been well-studied, and algorithms based on the nearest-neighbor model of base stacking [8, 9] predicts minimum free energy structures and melting temperatures [10, 11] with reasonably good accuracy. In contrast, the kinetics of hybridization remain poorly understood, and to date no models or algorithms have been reported that accurately predict hybridization rate constants from sequence and reaction conditions (temperature, salinity). This knowledge deficiency has adversely impacted the research community by requiring either trial-and-error optimization of DNA primer and probe sequences for new genetic regions of interest, or brute-force use of thousands of DNA probes for target enrichment.

Predictive modeling of hybridization kinetics faces two main challenges. First, the hybridization of complementary sequences can follow many different pathways, rendering simple reaction models inaccurate for a large fraction of DNA sequences. It is not practical to construct a comprehensive model that considers every potential DNA hybridization mechanism, due to the large variety of possible DNA sequences. Second, there is a very limited number of DNA sequences whose kinetics have been carefully directly, either in bulk solution [12–14] or at the single-molecule level [15–17]. One reason for the relative lack of data is the requirement of fluorophore-functionalized DNA oligonucleotides, which at roughly $200 per sequence becomes cost-prohibitive for the hundreds of experiments needed to establish sequence generality.

Here, we present a new approach to predicting DNA hybridization rate constants from sequence, which we call Weighted Neighbor Voting (WNV). WNV is inspired by machine learning concepts such as weighted k-nearest neighbor classification [18] and kernel smoothing [19]. In brief, each hybridization reaction (comprising a target sequence, a probe sequence, and a set of temperature and buffer conditions) is mapped to a set of feature values (Fig. 1). Features are scalar metrics that we rationally designed based on our understanding of potential factors that may influence hybridization kinetics. Each hybridization reaction is thus represented by a point in a high-dimensional feature space. Given a set of properly designed and weighted features, two hybridization reactions that are close in feature space are expected to exhibit similar kinetics. The rate constant of an unknown hybridization reaction is predicted based on the weighted average of observed rate constants for all experimentally tested reactions, with weights dropping exponentially for reactions that are farther away in feature space.

**FIG. 1:**
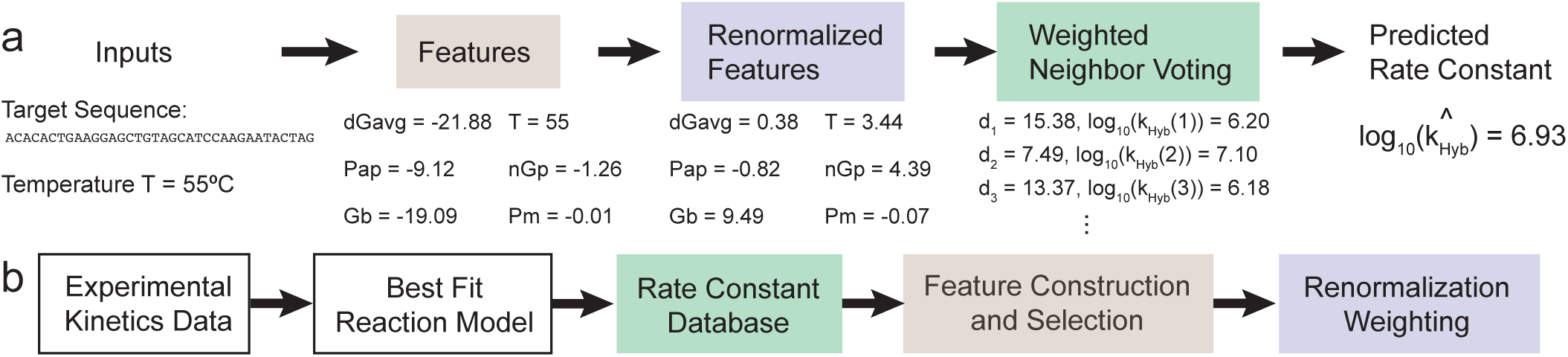
Method overview. **(a)** Rate constant prediction workflow for a user-defined target sequence and temperature. Six features are numerically computed from the input sequence and temperature, and subsequently renormalized. The target’s features values are compared to those in our database, and the observed rate constants in the database are integrated via a weighted voting system, with weight decreasing exponentially based on distance to the target in feature space. **(b)** Steps taken to generate final hybridization rate constant prediction model. Box colors mark the corresponding steps where the information is used in panel (a).

To create a sufficiently representative and sequence-general dataset for optimizing and validating features and weights, we experimentally characterized the kinetics of 210 individual hybridization reactions on 100 different pairs of complementary sequences. We rationally designed an initial pool of 38 features, which were pruned down by computational optimization to 6 in our final model. Under leave-one-out (LOO) cross-validation, our final WNV model predicts rate constants to within a factor of 2 for 74% of reactions, and within a factor of 3 for 92%. Next-generation NGS studies show a significant correlation *(R*^2^ ≈ 0.6) between the rate constants of DNA hybridization in single-plex vs. multiplex, suggesting that the current work is a good starting point for rational design and selection of DNA probes for highly multiplexed applications, such as target enrichment from genomic DNA.

## Experimental Results

To systematically but economically characterize the hybridization kinetics of many different sequences, we used the X-Probe architecture [20] that employ universal fluorophore and quencher-labeled oligonucleotides (Fig. 2a). Universal sequences (blue in Fig. 2a) were appended to the 5′ end of each target and the 3′ of each probe; the universal sequence for targets differs from that for probes. A universal fluorophore-labeled oligonucleotide was pre-hybridized to the probe, and a universal quencher-labeled oligonucleotide was pre-hybridized to the target. When the target and the probe solutions were mixed, the solution fluorescence was initially high because the fluorophore was delocalized from the quencher, but dropped over time as the hybridization reaction proceeds. The solution fluorescence at any given time can thus be linearly mapped to the instantaneous hybridization reaction yield.

**FIG. 2:**
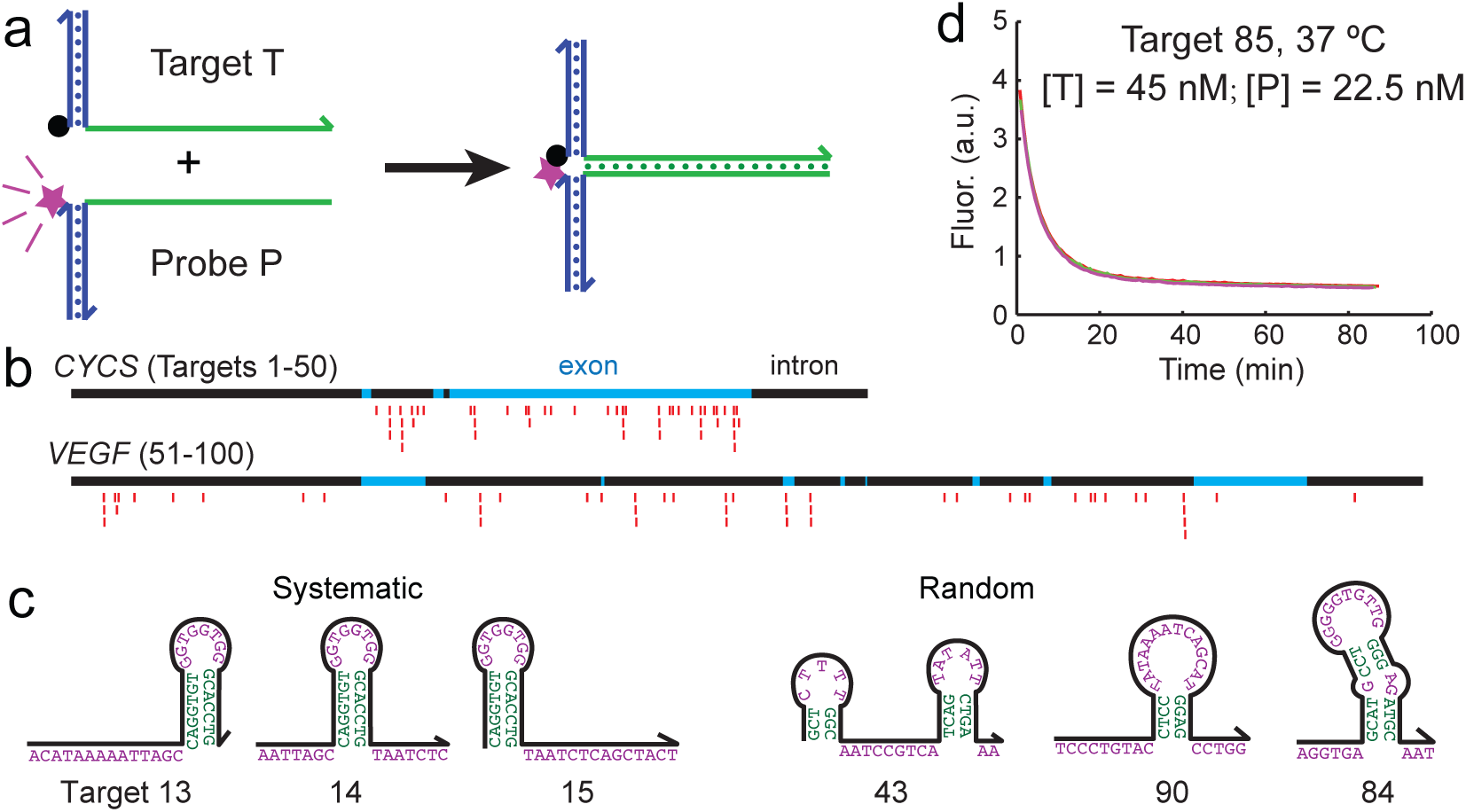
Experimental characterization of hybridization kinetics. **(a)** Fluorescent probes with universal functionalized oligonucleotides. The blue regions show universal sequences, and the green regions show the variable regions corresponding to the target or probe sequence. Fluorescence is initially high, and decreases as the hybridization reaction proceeds because the fluorophore (purple star) becomes localized to the quencher (black dot). **(b)** 100 different subsequences of the CYCS and VEGF genes were selected to be the target sequences. In this study, all target and probe sequences are 36 nt long (excluding universal regions). 25 targets for each gene were chosen randomly with uniform distribution across the entire intron and exon region, and the other 25 targets were selected as close overlapping frames to systematically test the position effects of secondary structures. **(c)** Examples of secondary structures encountered in target sequences. Shown are predicted minimum free energy (mfe) structures predicted for the target sequences at 37 °C. See Supplementary Table ST1 for sequences of the 100 targets. **(d)** Example kinetic traces (triplicate) of a hybridization reaction. All reactions proceeded in 5x PBS buffer. See Supplementary Section S1 for fluorescence traces for all 210 experiments.

To obtain a reasonable sampling of potential biological target sequences, we selected as targets 100 subsequences of the *CYCS* and *VEGF* genes, each target subsequence being 36 nucleotides (nt) long. Of the 50 targets for each gene, 25 of them were selected randomly with uniform position distribution across the gene, and the other 25 were selected systematically so that the effects of secondary structure position could be examined (Fig. 2b). For example, Fig. 2c shows likely secondary structures of 6 different targets at 37 °C; random target sequences exhibited a broader diversity of predicted secondary structures.

Fig. 2d shows triplicate kinetics traces for one hybridization reaction. A total of 210 hybridization experiments were characterized (100 reactions at 37°C, 96 at 55°C, 7 at 28°C, and 7 at 46°C); see Supplementary Section S1 for fluorescence traces for all experiments. There was very low experimental error in our fluorescence experiments; all triplicate data points agreed with each other to within 2%. To obtain maximally reliable experimental data for rate constant inference, we performed multiple experiments until determining a set of target and probe concentrations such that each hybridization reaction undergoes between 2 and 10 half-lives within the 80 to 180 minute observation time.

## Model Construction

### Hybridization rate constant (*k*_Hyb_) fitting

From the experimental kinetics traces, we wish to determine a single rate constant k_Hyb_ that describes the dominant hybridization kinetics pathway. However, a simple two-state T + P → TP reaction model fails to reasonably capture a significant portion of the kinetic behavior. Most notably, over 40% of the observed reactions appeared to asymptote to a final reaction yield of less than 85%, based on positive and negative control fluorescence values. Consequently, we considered 3 slightly more complex reaction models of hybridization, in order to evaluate which best fits the observed data (Fig. 3a).

**FIG. 3:**
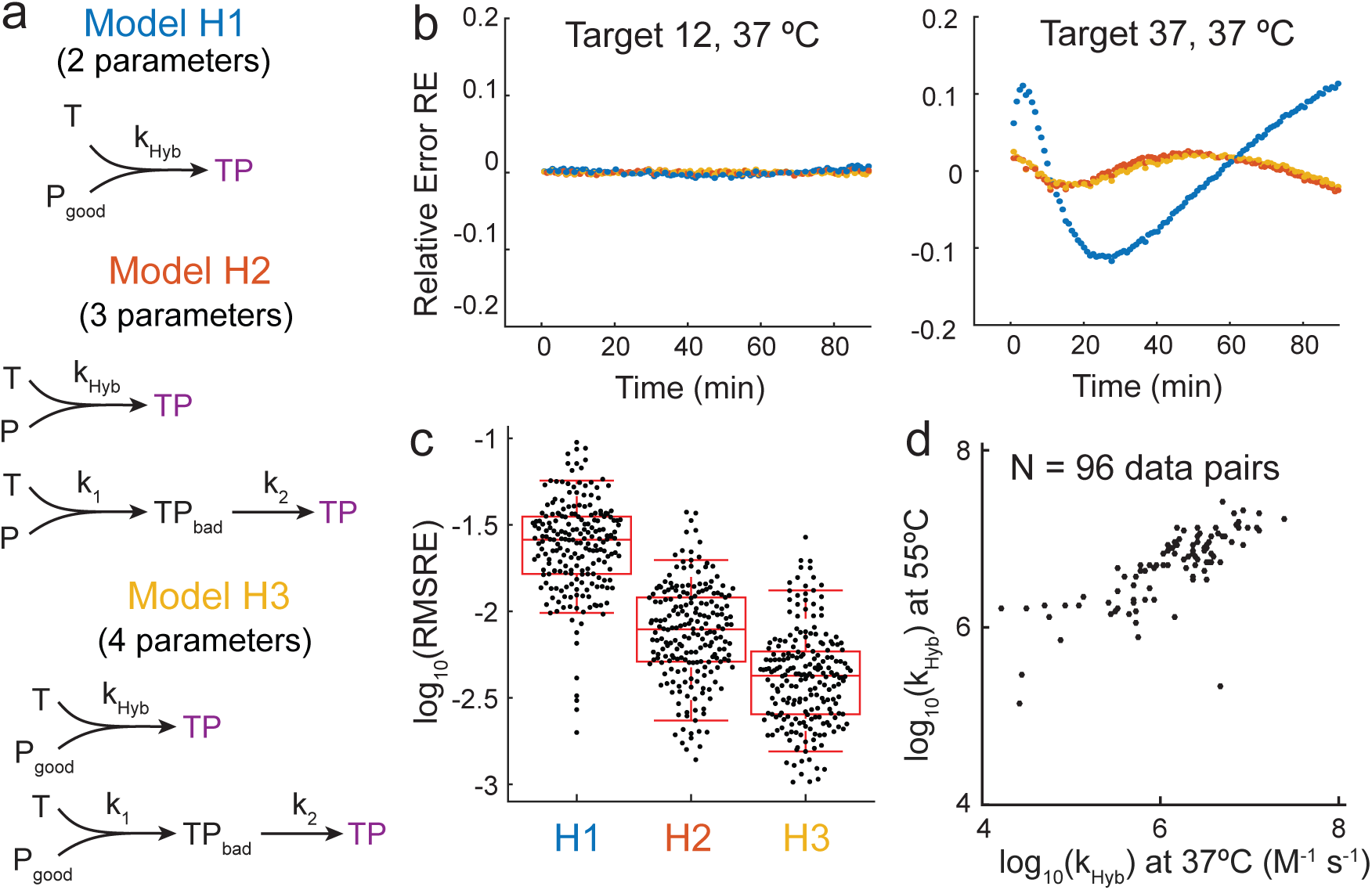
Hybridization model and rate constant parameterization. **(a)** Three different reaction models considered for fitting rate constant *k*_Hyb_ to fluorescence kinetics data. Model H1 is the simplest model and has 2 free parameters: *k*_Hyb_ and [P_good_]. [P_good_] denotes the concentration of properly synthesized probes that are capable of hybridization, with the remainder of the [P] assumed to be unhybridizable and constitutively fluorescent. Model H2 has 3 free parameters (*k*_Hyb_, *k*_1_, and *k*_2_), and Model H3 has 4 (*k*_Hyb_, [P_good_], *k*_1_, and *k*_2_). An even simpler model H0 with only *k*_Hyb_ as a parameter fails to reasonably fit the observed fluorescence data. **(b)** Two examples of fit quality for the 3 reaction models. The y-axis plots the relative error RE as a function of time for each model using best-fit parameters. For Target 12, all three models show low relative error across all time points, but H1 is significantly worse than H2 and H3 for target and 37. **(c)** Summary of fit performance for the three models across all 210 fluorescence kinetics experiments. Each point corresponds to the root mean square relative error (RMSRE) of all time points for a particular fluorescence experiment. Based on this result, we chose to proceed with H3 and its best-fit *k*_Hyb_ values for all subsequent studies. **(d)** Observed rate constants for 96 targets at 37 °C and 55 °C. 4 of the targets did not stably bind to their corresponds probes at 55 °C due to being A/T rich, and were not included in this plot. See Supplementary Section S1 for best-fit simulations using Model H3.

Model H1 assumes that the T + P → TP reaction is correct, but that a fraction of the probes P are poorly synthesized, or otherwise incapable of proper hybridization with target T or the accompanying fluorescence quenching. Thus, in addition to *k*_Hyb_, H1 has one extra fitting parameter: [P_good_]_0_, the initial concentration (or fraction) of viable probe P.

Model H2, in contrast, assumes that all probe P is correctly synthesized, but that some fraction of the T + P reaction undergoes an alternative pathway with rate constant *k*_1_ to result in a state TP_bad_ with high fluorescence. This frustrated state TP_bad_ may represent states in which T and P are co-localized by misaligned base pairs. Model H2 assumes that TP_bad_ undergoes first-order rearrangement with rate constant *k*_2_ to form the correct product TP. H2 has a total of 3 fitting parameters: *k*_Hyb_, *k*_1_, and *k*_2_. Model H3 is a simple combination of models H1 and H2, wherein there exists both a fraction of poorly synthesized P as well as the alternative pathway involving TP_bad_, and has a total of 4 fitting parameters: *k*_Hyb_, [P_good_]_0_, *k*_1_, and *k*_2_.

For each of our 210 fluorescence kinetics experiments, we used a custom stochastic fitting function to determine the best-fit values of each rate constant parameter for each model. Here, best-fit is determined as the minimal sumof-square relative error RE, where 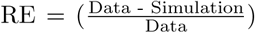. Minimum and maximum fluorescence values corresponding to 0% and 100% yields were determined through separate control experiments. Fig. 3b shows RE values of best-fit parameters for two hybridization reactions. While all three models describe the observed fluorescence data well for some reactions, other reactions show a significant difference among the three models. See Supplementary Section S1 for relative error plots for all kinetics experiments.

For each hybridization reaction, we have between 60 and 180 RE values, each corresponding to a time point at which fluorescence was measured. The RE values of each hybridization experiment are summarized as a single root mean square relative error (RMSRE) value, defined as

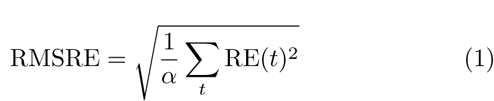

where *α* is the total number of time points *t* during which fluorescence was measured for the reaction. Fig. 3c shows the distribution of RMSRE values for each hybridization model; H3 appears to give the best overall fit to the data. We also evaluated more complex reaction models with additional fitting parameters, but these did not significantly improve RMSRE over H3 (data not shown). Consequently, H3’s best-fit *k*_Hyb_ rate constants were used for all subsequent work.

Fig. 3d summarizes H3’s best-fit *k*_Hyb_ values for paired hybridization experiments at 37 °C and 55 °C. The values of *k*_Hyb_ ranged roughly 3 orders of magnitude at both temperatures. Even among relatively fast reactions corresponding to target/probe sequences with relatively low secondary structure, there is still significant variation in hybridization kinetics. The large diversity of *k*_Hyb_ values for different sequences and the imperfect correlation between rate constants for the same sequence at different temperatures emphasizes the difficulty and need for a predictive kinetics model.

### Weighted Neighbor Voting (WNV) Model

To predict the rate constant of a new hybridization reaction, the WNV model checks the reaction for similarity against labeled instances (hybridization reactions with known rate constants) in our existing database, and allow each instance in the database to make a weighted “vote.” Instances that are more similar to the new reaction are weighted more heavily.

To quantitate the similarity or dissimilarity between two hybridization reactions, we abstract each reaction into a number of features. The value of each feature for a particular hybridization reaction is computable based on the sequences of the target and probe, and the reaction temperature and buffer conditions. Each hybridization reaction is thus a point in feature space. With an optimally designed and weighted set of features, the two points close in feature space should exhibit similar *k*_Hyb_ values. The converse is not necessarily true: two hybridization reactions with coincidentally similar *k*_Hyb_ values may possess very different feature values.

Mapping the hybridization reactions into feature space is important because targets that are similar in sequence space may not have similar hybridization kinetics, and vice versa, due to the sensitivity of secondary structure to small changes in DNA sequence in certain regions, but not in others. For example, oligonucleotide (2) with sequence “ACACACACTTAAAATTGTGTGTGTCCC” has higher Hamming distance to oligo (1) with sequence “ACACACACTTTTTTTTGTGTGTGTCCC” than oligo (3) with sequence “ACTCAGACTTTTTTTTGTGTGTGTCCC”, but is expected to exhibit much more similar kinetics in hybridization to each’s respective complement. In this case, one possible feature could be the number of base pairs formed in the hairpin stem of the minimum free energy structure: oligos (1) and (2) would have feature value 8, while oligo (3) would have feature value 6.

There are many potential approaches to the prediction of an analog desired parameter (*k*_Hyb_ in this paper) based on a set of features, the simplest of which is multilinear regression (MLR). We opted for WNV because WNV significantly outperforms MLR when the relationships between the desired parameter and the features are nonlinear. Simultaneously, WNV is a highly scalable framework, in the sense that additional labeled instances can easily be incorporated for improved prediction accuracy without requiring reoptimization of model parameters (feature weights). See Supplementary Section S2 for two simple examples comparing WNV and MLR.

### Feature Construction and Normalization

We started by rationally designing 38 potential features, each based on some aspect of DNA biophysics that we believed may influence kinetics. Supplementary Section S3 shows our full feature list and descriptions, as well as the correlation of each feature value with *k*_Hyb_; see Supplementary Table ST1 for values of the features for the 210 hybridization experiments. Fig. 4a shows Gb, one of the 6 features used in our final model; Gb can be thought of as the weighted average of the Δ*G*° of formation of the TP complex, based on probabilities of state existence/occupancy at equilibrium. Fig. 4b shows the relationship between the observed *k*_Hyb_ (according to model H3) and the value of Gb. There is significant correlation between *k*_Hyb_ and Gb; simultaneously, the relationship is not clearly linear. There may not be good physical interpretations of all effective features—in these cases, the feature in question is likely correlated with a yet-undiscovered complex feature with a firm physical basis.

**FIG. 4:**
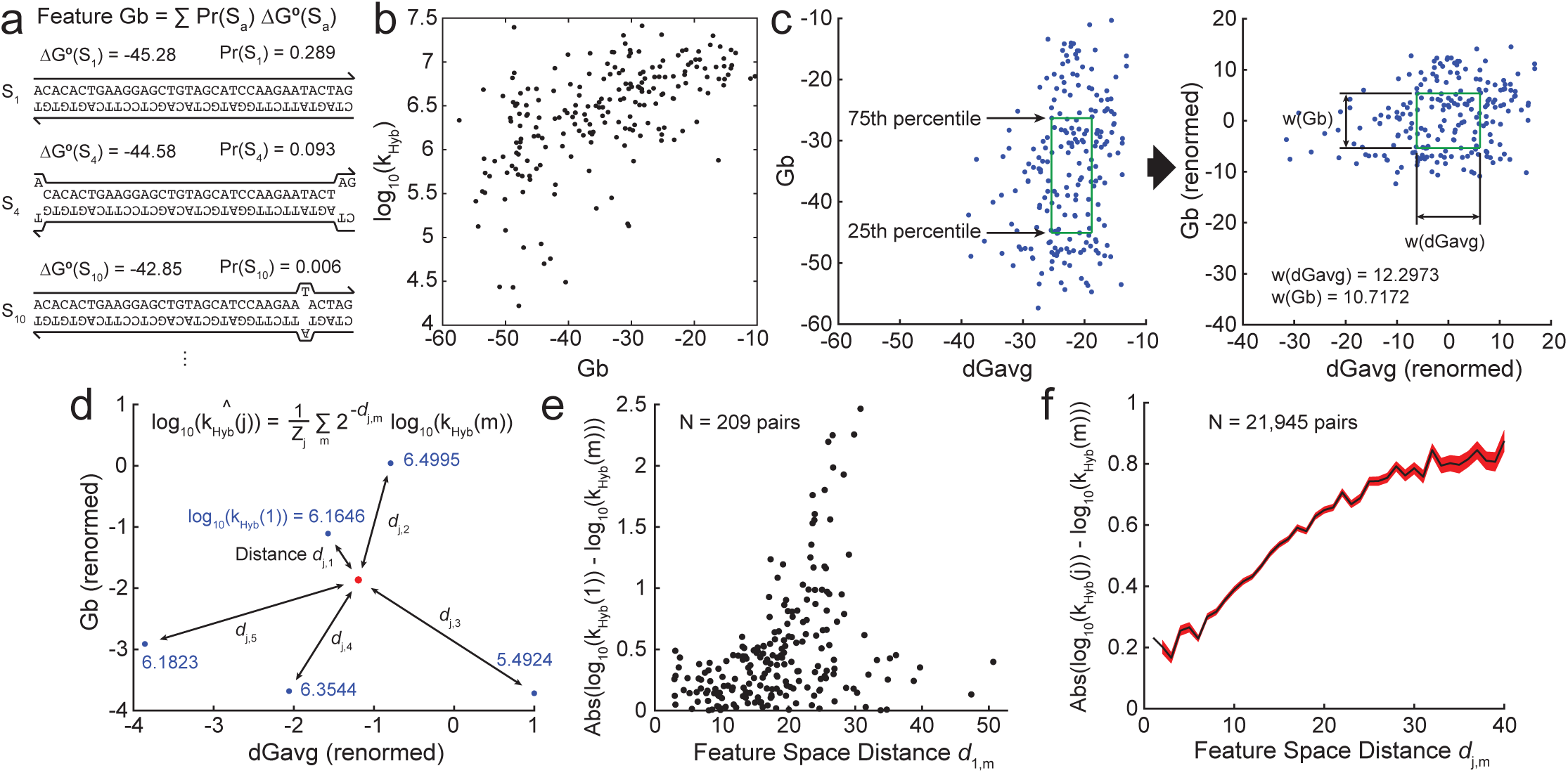
Rate constant prediction using the Weighted Neighbor Voting (WNV) model. **(a)** The values of a number of rationally designed features are computed based on the sequences of the target and probe, as well as the reaction conditions (temperature, salinity). Shown here is an example calculation for feature Gb, the weighted average Δ*G*° of the hybridized complex. The Δ*G*° of formation for a number of likely states S_*a*_ are computed using Nupack [11], and subsequently used to compute their probability of existence at equilibrium Pr(S_*a*_). **(b)** Relationship between the base-10 logarithm of the experimental hybridization rate constants *k*_Hyb_ (based on reaction model H3) vs. Gb values for the 210 hybridization experiments. There is moderate correlation between *k*_Hyb_ and Gb, indicating that Gb may be an effective feature for rate constant prediction. **(c)** Feature nenormalization based on 75th and 25th percentile values. Plotted to the left are the raw values of the Gb and dGavg features for all 210 experiments. These feature values are linearly transformed (re-normalized) based on a set of feature weights *w*(*i*): The 75th percentile value of a feature *i* is renormalized to 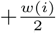 and the 25th percentile value is renormalized to *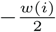.* **(d)** Given a reaction whose rate constant *k*_Hyb_ is to be predicted (red dot), the reaction’s renormalized feature values are first calculated and compared to the feature values of reactions with known *k*_Hyb_ values (blue dots). The feature space distance *d_j,m_* between the unknown reaction *j* and each known reaction *m* is used to determine the prediction weight of reaction *m.* Prediction weight drops exponentially with distance and is calculated as *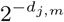*; *Z_j_* is the sum of all prediction weights involving *j.* Both the predicted rate constant 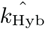 and the known rate constants are expressed in logarithm base 10. **(e)** Relationship between feature space distance *d* and the absolute value of difference in experimental rate constants (log 10) for two hybridization reactions. Here, feature space distances are calculated using the final 6-feature model (see Fig. 5), for one reaction (arbitrarily assigned *j* = 1) vs. all 209 other reactions. Pairs of reactions with small *d* generally have similar rate constants. The converse statement is not true because two very different reactions may coincidentally have similar rate constants. **(f)** Summary plot of Abs(log_10_(*k*_Hyb_(j)) - log_10_(*k*_Hyb_(m))) vs. feature distance *d_j,m_* for all 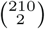 pairs of experiments. The black line shows the mean, and the red region shows ±1 standard deviation on the mean.

The features that we constructed had different units and different ranges of values. In order to calculate a distance between two hybridization reactions, it is necessary to normalize the different features into a consistent scale. Because the distributions of most feature values were distinctively non-Gaussian for our 210 reactions, normalization was performed based on the interquartile range: the 75th percentile feature value is mapped to a score of 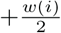, and the 25th percentile value is mapped to *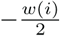* (Fig. 4c), where *w*(*i*) is the weight of feature *i.* The feature space distance *d_j,m_* between an unknown reaction *j* and a known reaction *m* is calculated as a Euclidean distance:

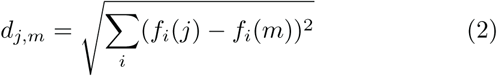

where *f_i_*(*j*) is the value of renormalized feature *i* for reaction *j* (Fig. 4d). Because a feature *i* with larger weight *w*(*i*) allows a larger range of scores, it can contribute more to the distance between two hybridization reactions. Fig. 4ef confirms that the difference in *k*_Hyb_ values for a pair of reactions increases with feature space distance *d.*

### Rate Constant Prediction

From a database of hybridization experiments *m* with known *k*_Hyb_(*m*) and renormalized feature values, our WNV model makes the following prediction for *k*_Hyb_(*j*) of an unknown hybridization reaction *j*:

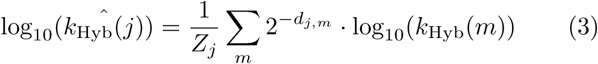

where 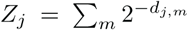 is the “partition function” of the distances involving reaction *j*.

To quantitate the overall performance of a particular WNV model (defined by its set of features and corresponding feature weights *w*(*i*)), we constructed the following “Badness” metric:

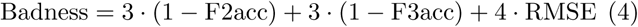

where F2acc is the fraction of all predicted reactions *j* in which predicted 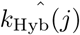 and the experimental *k*_Hyb_(*j*) agrees to within a factor of 2, F3acc the fraction that agrees to within a factor of 3, and

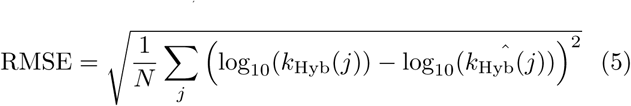

is the root mean square error of the logarithm of the hybridization rate constant (where *N* = 210 is the number of experiments).

We chose to use this Badness metric rather than RMSE only (i.e. a least-squares fit) because we felt that it is more relevant for many applications involving the design of DNA oligonucleotide probes and primers: Rather than marginally improving the predictions of outlier sequences that are off by more than an order of magnitude, our Badness metric emphasizes instead improving the fraction of predictions that are correct to within a factor of 3, or better yet within a factor of 2. Simultaneously, to allow efficient computational optimization of feature weights, the Badness metric to be minimized cannot be locally flat, so RMSE is included as a component of Badness. Use of different Badness metrics will result in optimized feature weights that exhibit a different tradeoff between the magnitude and frequency of large prediction errors.

### Feature Selection and Weighting

All 38 potential features we constructed showed significant correlation with *k*_Hyb_, but it is inappropriate to include all of these in our WNV prediction model both because several features may consider redundant information, and because large sets of feature weights are computationally difficult to optimize. We first manually pruned the list of potential features down to 17 most promising features, based on single-feature WNV performance (using each feature’s optimized feature weight). Due to the complexity and nonlinearity of the Badness landscape over the feature weight parameter space, it was not feasible to determine an analytic solution of optimal weights. Instead, we used a stochastic numerical optimization algorithm to find weight values that achieve Badness minima.

Next, we implemented a greedy algorithm in which individual features that best improve the Badness at each round are iteratively added to an initially empty feature set. Fig. 5a shows that the Badness decreases as the number of features included increases up to 8; at 9 features, the WNV model showed no additional Badness improvement. Also plotted in Fig. 5a are the Badness of for MLR model using various numbers of features. Up to 4 features, the WNV and MLR models provide similar prediction accuracies; however, WNV continues improving with additional features whereas MLR performance plateaus. See Supplementary Section S4 for MLR optimization details.

**FIG. 5:**
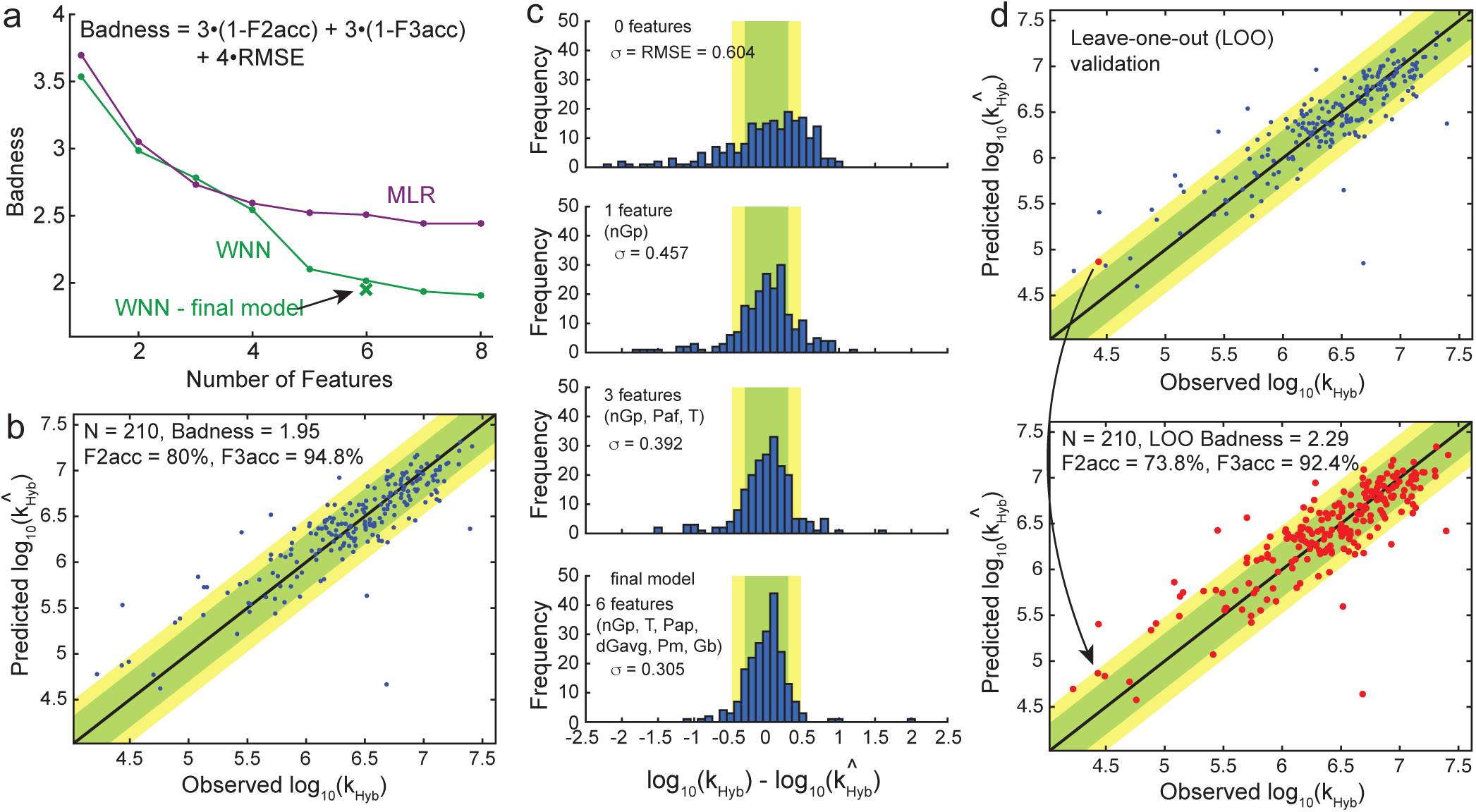
Prediction accuracy of the WNV model. **(a)** The green dots show the results of a greedy optimization algorithm in which individual weight-optimized features are sequentially added to the WNV model. The Badness metric (y-axis) is calculated based on the fraction accurate to within a factor of 2 (F2acc), fraction accurate to within a factor of 3 (F3acc), and the root-mean-square error (RMSE). The final 6-feature WNV model (green X) comprise the 6 features from the 8-feature model with the largest weights. Performance of a multilinear regression (MLR) model is also shown for comparison; see Supplementary Section S4 for MLR feature selection and weight optimization details. **(b)** Rate constant prediction performance of the final WNV model, using feature weights were optimized on all 210 experimental reactions. Each blue dot corresponds to the prediction of each reaction’s rate constant based on the data from the other 209 reactions’ rate constants. **(c)** Histograms of experimental minus predicted log_10_(k_Hyb_) values for WNV using 0, 1, 3, and 6 features. The 0-feature model represents a naive model predicting the same log_10_(k_Hyb_) value for all sequences, and the 1-feature model represents a simple model that considers only secondary structure. **(d)** To estimate the accuracy of the final WNV model on prediction of rate constants for new sequences, we performed leave-one-out (LOO) validation. In the top panel, feature weights were optimized on the 209 blue data points, and this model was used to predict the rate constant of the single “unknown” red data point. The bottom panel summarizes the prediction performance of the 210 distinct models (with weights optimized on different sets of 209 points).

The optimized feature weights for the 8-feature WNV model included two features very small weights (*w <* 0.1); these were removed, and the final WNV model consist of the following 6 features: dGavg, Pap, Gb, T, nGp, and Pm, with weights of 12.30, 11.89, 10.72, 6.88, 6.54, and 0.94, respectively. A brief text description of these each feature follows; see Supplementary Section S3 for additional feature details. dGavg corresponds to the sum of the Δ*G*° of binding for all subsequences of the target weighted by the probability of all nucleotides of the subsequence being unpaired. Pap corresponds to the sum of the probability-weighted Δ*G*° of the strongest continuous subsequence that is expected to be unpaired. Gb was described in Fig. 4a and corresponds to the sum of the probability-weighted Δ*G*° of formation of the target-probe complex. T is the reaction temperature in Celsius. nGp corresponds to the partition function energy of the probe secondary structure, as calculated by Nupack. Pm is similar to Pap, but is calculated for misaligned target-probe complexes.

Fig. 5b shows the accuracy of the final 6-feature WNV model. Each blue dot plots the predicted 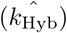 value vs. the experimentally observed log_10_ (k_Hyb_) value for a single hybridization experiment. Each prediction was performed using 209 labeled instances (all reactions except the one to be predicted), using the feature weights trained on all 210 data points. Predictions were accurate to within a factor of 2 for 80% of the reactions, and within a factor of 3 for 94.8% of the reactions. For comparison, Fig. 5c shows histograms of the distribution of prediction errors in log_10_ (k_Hyb_) for WNV models using 0, 1, 3, and 6 features.

### Leave-one-out Validation of Final WNV Model

The fact that the final model’s feature weights were fitted to all 210 experiments raises potential concern regarding whether the WNV model’s prediction accuracy would generalize to new hybridization reactions, because the latter’s (unknown) rate constant cannot be used for training feature weights. Here, we performed leave-one-out (LOO) validation on the model to study the generalizability of the WNV model.

In our LOO studies, we performed 210 separate feature weight optimizations, each using a different set of 209 hybridization experiments. Thus, each of the 210 models possessed different feature weights, and each model was used to predict the hybridization rate constants of the single hybridization experiment not included for its feature weight optimization (red dot in top panel of Fig. 5d). The aggregate performance of these 210 LOO models are shown in the bottom panel of Fig. 5d. F2acc and F3acc are marginally lower than in Fig. 5b at 73.8% and 92.4%, respectively.

To help the research community predict hybridization rate constants for DNA oligo probes and primers, we have constructed a web-based software tool, available at http://nablab.rice.edu/nabtools/kinetics The software typically completes predicting *k*_Hyb_ within 30 seconds, with the bulk of the computing time devoted to computation of the Pap and Pm feature values. It is currently seeded with the 210 hybridization experiment results performed in this paper, but will be updated with additional hybridization experiment results in the future, which should further improve prediction accuracy.

### Enrichment from Human Genomic DNA

The human genome is over 3 billion nucleotides long, but the coding regions that form the exome collectively only span 30 million nucleotides, or 1% of the genome. Within the 20,000 genes of the exome, typically there are only between 10-400 are that are relevant to any particular disease. Consequently, solid-phase enrichment of relevant gene regions using highly multiplexed hybridization of synthetic DNA oligonucleotide probes [6] is the preferred approach for targeted sequencing.

Current commercial multiplex hybrid-capture panels generally use a very large number of synthetic probe oligonucleotides to fully tile or overlap-tile the genomic regions of interest; for example, the whole exome requires more than 200,000 distinct oligonucleotide probe species. Due to the large number of oligo species involved, the concentration of each species is thus necessarily quite low (tens of picomolar), resulting in hybrid-capture protocols that typically span at least 4 hours, and more frequently more than 16 hours. Because of the varying hybridization kinetics of different probes (Fig. 3d), it is likely that many probes do not contribute significantly to hybridization yield, and in fact slow down the hybrid-capture process by forcing lower concentrations of the fast-hybridizing probes.

To experimentally test this possibility, we first applied our hybridization rate constant prediction algorithm to all possible 36 nt probes to exon regions of 21 genes. Because the exon regions are typically 3000 nt long, this corresponds to roughly 3000 possible probes. Predicted rate constants typically range about 2 orders of magnitude (see Supplementary Fig. S5-4), with the fast (≥95th percentile) probes being typically a factor of 3 faster than median probes (≈50th percentile). NGS hybrid-capture enrichment typically uses probes longer than 36 nt (e.g. Agilent SureSelect uses 120 nt probes), but there is likely a similar if not greater range of hybridization kinetics rate constants for longer probes due to the greater possibility of secondary structure and nonspecific interactions.

Subsequently, we picked a total of 65 fast probes and 65 median probes across the exon regions of 21 different cancer-related genes. The expectation is that after a 24 hour hybridization protocol, the fast and median probes would produce similar reads, but with a short 20 minute hybridization protocol, the fast probes would exhibit significantly greater reads than median probes (Fig. 6a). Our library preparation protocol is summarized in Fig. 6b; all 130 probes are hybridized to the adaptor-ligated DNA simultaneously. However, the number of reads aligned to a particular probe is not directly proportional to its hybridization yield, due to well-documented sequencing bias [24, 25]. For example, some adaptor-ligated amplicons exhibit significant secondary structure and is less efficiently PCR amplified during normalization, or less efficiently sequenced due to lower flow cell binding efficiency. For this reason, 15 fast and 15 median probes targeting 4 genes resulted in less than 100x sequencing depth, and were excluded from subsequent analysis (see Supplementary Section S5); we do not believe this to affect the conclusions from our genomic DNA enrichment study.

**FIG. 6:**
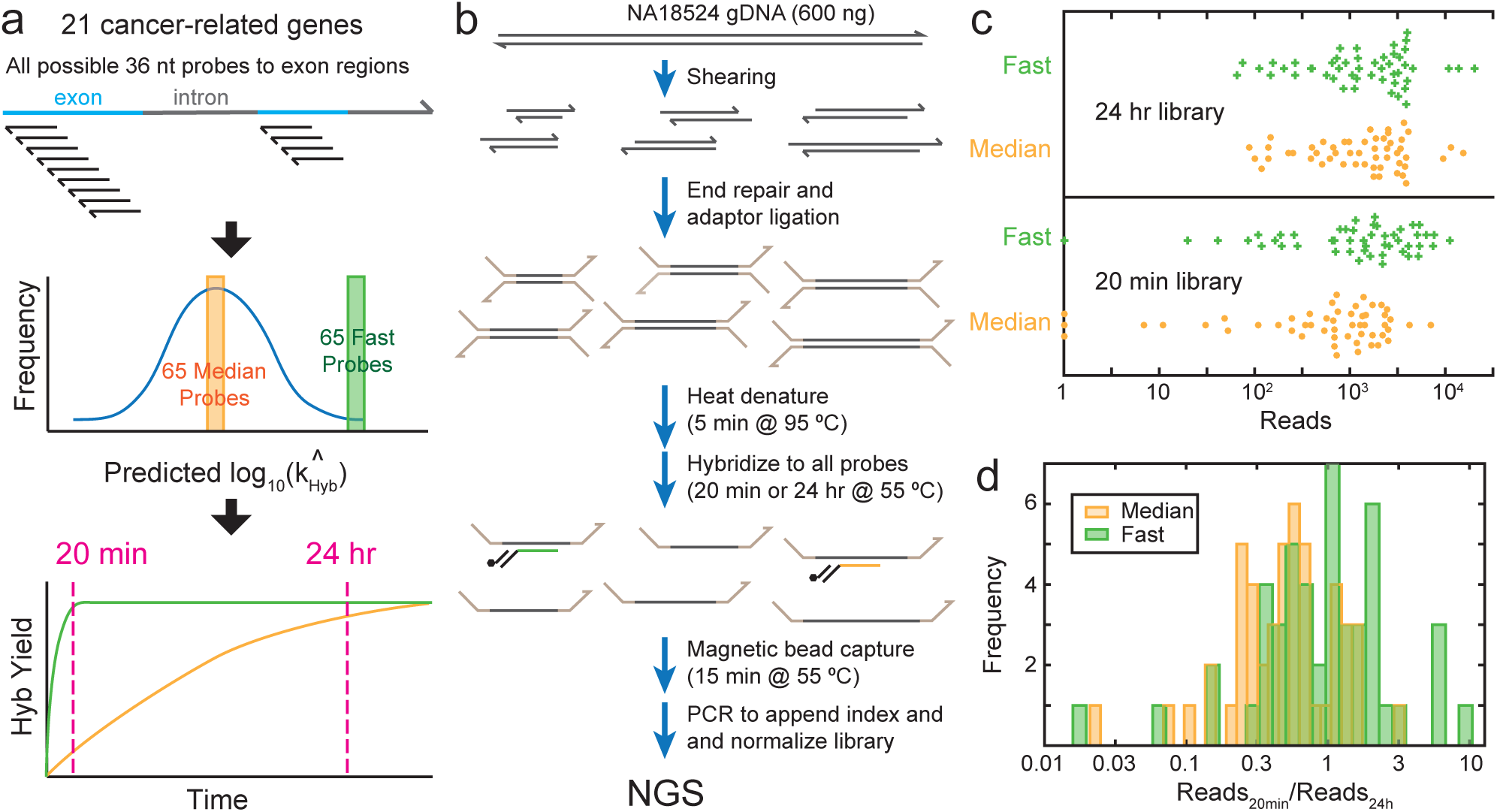
Comparison of probes predicted to possess median vs. fast hybridization kinetics for enrichment from human genomic DNA. **(a)** Hybridization rate constant k_Hyb_ were predicted for all possible 36-mer hybridization probes to the exon regions of 21 cancer-related genes. The middle and lower panels express the idea behind probe selection and library design, and do not accurately reflect kinetics distributions or trajectories of any particular gene or probe; see Supplementary Fig. S5-4 for the distribution of predicted k_Hyb_ for the AQP1 gene. **(b)** Genomic DNA enrichment and library preparation workflow. All hybridization probes were present at 50 pM concentration. See Methods for detailed protocol. **(c)** Beeswarm plot of NGS reads aligned to each probe, excluding 15 fast and 15 median probes to 4 genes with low read depth (see Supplementary Section S5). In the library in which probes were hybridized to the fragmented gDNA for 24 hours (top panel), there is no significant difference in the read count distribution between the median and fast probes. In the 20 minute hybridization library, the fast probes showed significantly higher reads than the median probes, indicating that the probes our algorithm predicted to be faster did in fact provide a higher degree of hybridization within 20 minutes. **(d)** Ratio of aligned reads in the 20 minute library to the 24 hour library for each probe. A high ratio indicates fast hybridization kinetics; ratio can exceed 1 because libraries were normalized, so that fast probes are more dominant and occupy more reads in the 20 minute library.

Our comparison of reads for the 20 minute hybridization library and for the 24 hour hybridization library indicates that the probes predicted to be fast on average exhibited both a 2-fold increase in reads in the 20 minute library, and a 2-fold increase in the ratio of reads at 20 min vs. 24 hours. This is slightly worse than our algorithm’s predicted 3-fold difference between median and fast probes, but understandable given that our rate constant prediction algorithm was trained on single-plex hybridization rather than on multiplex hybridization. Our calibration experiments (Supplementary Section S5) indicate that the correlation constant between single-plex and multiplex k_Hyb_ values are roughly *r*^2^ = 0.6.

Our results thus suggest that sparse hybrid-capture enrichment panels would produce faster kinetics at a significantly lower cost. Rather than fully tiling or overlap-tiling the genetic regions of interest, it would be better to use a higher concentration of a few probes with fastest hybridization kinetics. Multiple probes are only needed insofar as biological genomic DNA may be fragmented, and a different probe is needed to capture each fragment. With the notable exception of cell-free DNA [26], most genomic DNA from clinical samples are longer than 500 nucleotides.

The concentrations of the probes used for this study was 50 pM per probe, and was intentionally selected so as to be similar to the concentrations of probes used by commercial enrichment kit providers At 50 pM concentrations, up to 200,000 probes can be used and the total oligo concentration would still be at a reasonable 10 *μ*M. At the significantly (e.g. 10x) higher individual probe concentrations that become feasible with a sparse coverage of target genetic regions, even the 20 minutes allotted here for hybridization could be further reduced, greatly speeding up the NGS library preparation workflow from current practice of 4-24 hours.

## Discussion

In this work, we combined the rational design of features and the WNV framework with computational optimization of feature selection and feature weights, resulting in a final model that is capable of accurately predicting hybridization kinetics rate constants based on sequence and temperature information. The final WNV rate constant prediction model is highly scalable and easily incorporates new experimental data to provide improved predictions, without requiring model retraining. With every additional hybridization experiment and its accompanying fitted *k*_Hyb_ value, the 6-dimensional feature space becomes denser, ensuring that on average a new hybridization experiment will be closer to an existing labeled instance. Thus, prediction accuracy will further increase and as we and other researchers collect additional hybridization kinetics data.

To seed the model with a reliable initial database of labeled instances that is representative of the diversity of genomic DNA sequences, we experimentally characterized the kinetics of 210 hybridization experiments across 100 biological target sequences using fluorescence. The X-probe architecture allowed us to economically study kinetics for a reasonably large number of target sequences, but extra nucleotides of the universal arms may cause hybridization kinetics to differ slightly from that of a standard single-stranded probe. For example, there may be a systematic bias towards lower rate constants because of the reduced diffusion constants. Nonetheless, because all targets/probes use the same universal arm sequences, it is likely that the relative ordering of rate constants is preserved.

In this work, we started with 38 rationally designed features that we eventually pruned down to 6 in the final model. The high LOO validation accuracy of the WNV model indicates that these features capture a significant, if not majority, portion of the complexity of the hybridization process. Simultaneously, there remain pairs of experiments in our database with similar feature values (feature space distance *d* ≤3) but with 3-fold differences in *k*_Hyb_. This implies the existence of undiscovered features that would distinguish these pairs of experiments; additional insight and creativity from the community in designing additional features would be welcomed.

The hybridization reactions experimentally characterized in the work were all performed in 5x PBS buffer, and all target and probe sequences were 36 nt long. These experiment constraints were designed to reduce the diversity of hybridization reactions, in order to ease the training of the WNV model. We plan to expand experimental studies to these various conditions, in order to allow the WNV model to accurately account for buffer conditions and probe length. Additionally, with genomic DNA targets, the long-range secondary structure and the fragmentation pattern of genomic DNA targets should also be considered. An expanded model to accommodate varying length targets and probes (including targets overhangs) and other buffer conditions will require the construction of new features.

Multiplex hybrid-capture panels for enriching target regions from genomic DNA is commonly used in targeted sequencing for scientific and clinical studies. In the absence of reliable kinetics prediction software, researchers and companies have taken a brute-force probe design approach, using fully tiled or overlapping-tiled probes to cover genetic loci of interest. While this approach ensures the presence of at least some fast-binding probes, it is both expensive (in terms of synthesis and QC of thousands of probes) and results in slower workflows. Accurately predicting multiplexed hybridization kinetics will enable precision design of sparse, high-performance probe panels for target enrichment.

## Acknowledgements

The authors thank Sherry X. Chen for assistance with NGS sequence alignment. This work was funded by NIH grant R01HG008752 to DYZ.

## Author contributions

JXZ, LRW, and DYZ conceived the project. JXZ and AWZ performed the experiments. ND and AP performed hybridization reaction model fitting and selection. JXZ, JZF, BY, and RP performed feature construction. WD and DYZ performed WNV model construction and optimization. ND, BY, RP, and RP performed MLR model construction and optimization. DYZ wrote the manuscript with input from all authors.

## Additional information

Correspondence may be addressed to DYZ. There is a patent pending on X-probes used in this work. There is a patent pending on the WNV model of hybridization rate constant prediction.

